# Sequential development of embryoblast-like memory entities in human cancer tissues: An evolutionary self-repair structure with pluripotentiality

**DOI:** 10.1101/2020.10.02.324376

**Authors:** Jairo A Diaz, Liliana Sánchez, Luis A Diaz, Mauricio F Murillo, Laura Poveda, Oscar F Suescun, Laura Castro

**Author notes:** Address correspondence to: Jairo A Diaz MD Pathologist Hospital Departamental de Villavicencio Pathology Laboratory Villavicencio Meta Colombia. Contributed equally to this work (Jairo A Diaz, Liliana Sánchez, Luis A Diaz).

## Abstract

Hidden collective organization of cancer cells can partially or completely return to embryoid genotype-phenotype with the plasticity to transform their morphology on cell embryoblast-like memory entities by expression of dormant genes that arise from embryogenesis.

After hundreds of driver mutations, cancer cells gain new abilities or attributes and recapitulate early stages of embryogenesis. Our findings document how malignant tissues reactivated ancestral storage memory and elaborate inside tumor glands spiral- pyramidal-fractal chiral crystals (Tc) as geometric attractors proteins and biomimicry the primitive cellular blastocyst embryoblast fluid-filled cavity. The resultant evolutionary embryoblast-like entity has higher survivability and spatial cephalic-caudal growth organization with pluripotentiality that carry the correct DNA instructions to repair, and regenerate. The isolation and manipulation of these order structures can guide and control the regenerative pathway mechanism in human tumors as follows: modify and reprogram the phenotype of the tumor where these entities are generated, establish a reverse primordial microscopic mold to use the swirlonic collective behavior of cellular building blocks to regenerate injured tissues, convert cancer cells to a normal phenotype through regeneration using the organizational level and scale properties of reverse genetic guidance, global control of mitotic activity and morphogenetic movements avoiding their spread and metastasis, determining a better life prognosis for patients who incubate these entities in their tumors compared to those who do not express them. An emergent self-repair order structure, biological template to develop targeted therapeutic alternatives not only in cancer but also in treatment of autoimmune, viral diseases, and in regenerative medicine and rejuvenation.

## Introduction

The morphological patterns of malignancy have traditionally been exclusively based on the change in size and shape of cells or nuclei, abnormal mitosis and pleomorphic nuclei, hyperchromatic coarse clumped chromatin, a change in the nucleus/cytoplasm ratio, loss of polarity, and disordered growth. We propose that this approach to cancer is incomplete.

In 1908, a pathologist at the University of Manchester, Charles Powell White, noticed that there was something more than hyperchromatism and pleomorphism in the tumor tissues. White [1] observed the presence of crystals in malignant tumors based on the premise that alcohols dilute the crystals. The author relates these crystals in some way to the proliferative activity of the tumor, identifying them both in sarcomatous lesions and carcinomas. The crystals are made up of cholestyramine, and fatty acids seem to be associated with cell proliferation rather than with cell degeneration. In 1963, George Rose from the Biology Department of the University of Texas identified unusually large extracellular crystals and particles produced by **embryonic** chick cells in special tissue culture environments, including helical, tubular, ribbon-like, triangular, hexagonal, rhomboidal, and filamentous forms [2]. The origin of these forms was detailed biochemically, but their usefulness in the culture environments in which they were found remained obscure.

In 2007, in a preliminary study, we described and documented the self-assembly of geometric **triangular** chiral hexagon crystal complexes (GTCHCs) in human pathologic tissues at macroscopic and microscopic levels, particularly in cancer tissues [3,4]. The genesis of these complexes occurs through intercellular cancer collisions that lead to the degradation of membrane ejected actin filaments in the form of rotation–domain interactions – that is, pairs of filaments with left- and right-hand sub-patterns of spin spirals [5]. Recent observations confirm the previous findings on GTCHCs identified in cancer tissues 10 years ago: interfacial geometry dictates cancer cell tumorigenicity [6], and matrix geometry determines the optimal cancer cell migration strategy and modulates responses to interventions. Whereas tumor cells exploit geometry for metastasis [7], the geometry helps confined cells to acquire a stem cell phenotype [8]. Today, we propose, in accordance with our findings that this geometry of spirals and **pyramidal-triangular** patterns that we call triplet crystals (Tc) represents the crystalloid mold proteins on which the spatial order that gives rise to what we call embryoblast memory entities in malignant tumors is built.

Tumorigenesis resembles the self-organizing process of early embryo development. With the recent profound advances in the field of developmental biology, it has become apparent that the early development of embryos shares many similarities with cancer development in terms of both biological behaviors and similarities in the cell invasive epigenetic regulation of gene expression and protein profiling. Thus, it is evident that tumorigenesis mimics a self-organizing process of early embryo development [9–15]. The aim of this research is to demonstrate the intimate connection that cancer has with embryogenesis. **Cancer recapitulates the early stage of embryogenesis**

## Methods

We re-examined all of our materials, collected In the past five(5) years in which we identified recurrent patterns of pyramidal-triangular and spiral chiral crystals (Tc) as geometric attractors in cancer tissues. corresponding to more than 1077 microscopic/macroscopic specimens, including carcinomas, adenocarcinomas, and sarcomas.

### Neuron-specific enolase–immunostaining

This isoenzyme, a homodimer, is found in mature neurons and cells of neuronal origin. Detection of NSE with antibodies can be used to identify neuronal cells and cells with neuroendocrine differentiation.

Sixty formalin-fixed and paraffin-embedded tissue sections with the most representative hot spot of Tc identified in malignant tumors were analyzed using neuron-specific enolase. We performed immunohistochemistry using the standard protocol method with paraffin sections. The scoring was done as follows: Ni (no immunostaining); low (10% or less immunopositivity); or high (>10% immunoreactive cells).

This study was approved by the ethics subcommittee of the University Cooperative of Colombia, Villavicencio, Colombia, No 9108 /2016 and followed the guidelines of the Ministry of Health (No. 8430 of 1993) and the principles established by the Declaration of Helsinki. All patients signed an informed consent form for their biological materials to be used for diagnostic and research purposes.

It is important to mention that the images documented in this work belong to living patients who have a natural history of cancer, who have not previously undergone chemotherapy or radiotherapy. The size of the giant masses documented show that they have developed over approximately nine (9) years, with silent clinical growth. When these large masses were resected, none of these patients had developed metastases and it was for this specific reason that they were given the option of surgery as their condition allowed it

### Statistics analysis

#### Description of the statistical analysis

We have been extensively verifying and documenting for more than ten years that in a large number of tissues with cancer, that is, carcinomas, adenocarcinomas, and sarcomas, cells that assume a triangular and spiral pattern appear recurrently, which we know are generated from geometric chiral crystals that we call them Tc. We analyzed 1050 cases of cancer and calculated the presence of these geometric complexes in these tumors. Additionally, we selected 57 cases with the greatest representativeness of these geometric complexes and carried out immunohistochemical studies of Neuron-Specific Enolase, as a biomarker of neuronal differentiation, demonstrating that tumor cells that assume a geometric pattern acquire neuronal differentiation in turn.

A descriptive cross-sectional study was developed, a non-probabilistic sampling method was employed for convenience to select samples of malignant cancer tissues from prevalent cases.

We collected and re-examined the tissue samples looking for recurrent patterns of pyramidal-triangular and spiral chiral crystals (Tc) as geometric attractors to estimate the prevalence proportions (PP) as a *“point estimator”* for confidence interval of 95%. The data are presented in absolute and relative (%) frequencies tables and bar plots. The chi–squared (X2) probability distribution was employed for (n-1) degree freedom (df) and considering statistical significance with p≤0.05 for the null hypothesis the true probability is not equal to 0.5. Then, we randomly selected 60 samples of cancer tissues, for neuron-specific enolase–immunostaining, and estimated the (PP) with the same probability distribution mentioned earlier (X^2^). The R 4.1.2 study version was used for the statistical analysis.

## Results

1077 malignant cancer tissues samples were collected, the geometric complexes (Tc) were identified in 1050 cases; the prevalence proportion (PP) was 97.4% (IC 95% 0.965 - 0.984, X^2^ = 977.42, df = 1, p-value < 0.01); when we randomly selected 60 samples to neuron-specific – enolase was applied; we found a positive test in 57 samples, with a prevalence proportion (PP) of 95% (IC 95% 0.874 - 0.994, X^2^ = 50.417, df = 1, p-value < 0.01), see table 1 and Graph 1.

**Table 1.** Descriptive analysis.

| Variable | Absolute frequency (n) | Relative frequency (%) | Confidence interval 95% | Chi squared ($\chi^2$) | p - value |
| --- | --- | --- | --- | --- | --- |
| Geometric complexes (Tc) | 1050 | 97.4 | 0.965 - 0.984 | 977.42 | <0.01 |
| Neuron-specific enolase – immunostaining | 57 | 95 | 0.874 - 0.994 | 50.417 | <0.01 |

### Photomicrography evidence

We were able to capture a unique and perhaps unrepeatable image: the evolutionary cycle of structures with an embryoblast-like phenotype generated from squamous epithelial cells injured by Human Papilloma Virus in a Uterine cervical cytology with corresponding biopsy confirmation **of** cancer in situ. The cycle consists of 12 stages that show step by step how these structures are gestated.

As can be seen in the cervical smear images, these entities are gestated from fractal crystalloid protein memory modules that have an **embryoblast-like phenotype,** where clusters of malignant cells are aligned in progressive order sequences via rotating vortices. This mechanism is activated when the cell suffers irreversible damage from specific mutations, as in this case through the action of the Human Papilloma Virus (Fig 1).

**Figure 1.**
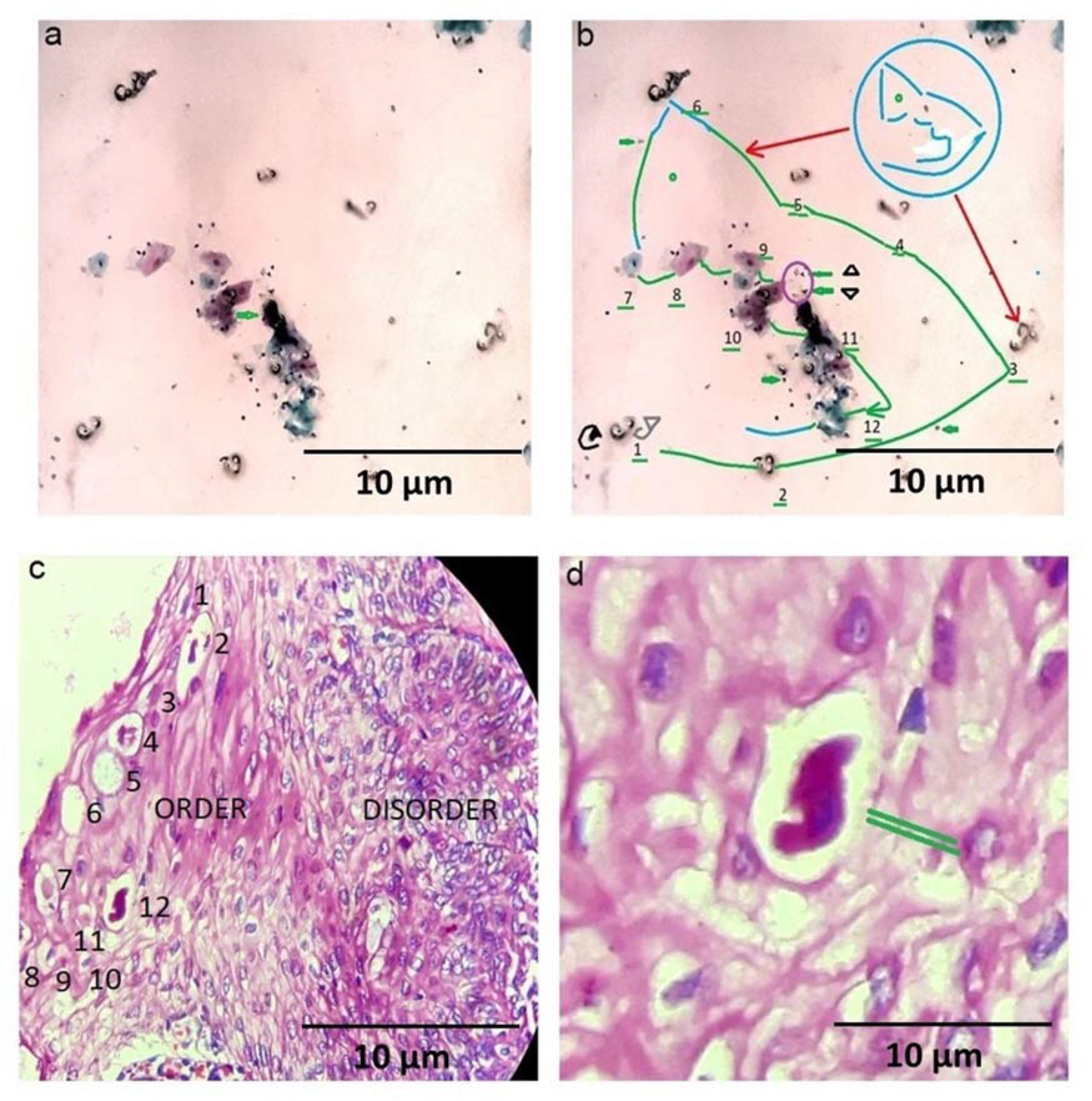
Self-repair structure: Cytologic smear and biopsy illustrates clusters of malignant cells aligned in progressive order sequences generating embryoblast-like entities being generated by spiral rotating vortices orientated in opposite directions. Swirlonic state observe the interface between crystal and cells intimately correlated with spatial-cephalic-caudal growth organization

We identified in the metamorphosis of these entities three memory modules

#### a) Protein Crystalloid memory module

Stages 1 to 6 (Fig 1).

We can observe in fig 1 a-b, cervical smear, the intelligent sequence of a chain of events. Protein *fractal* crystalloid entities measuring 4 to 9 ums, appear separate, aligned, and functionally interconnected. In stage 1 we can observe a polar entity made up of two (2) molecular crystals proteins of triangular and spiral shapes that appear in a **chiral** position. We had previously described this event as GTCHC complexes [3] With this evidence, we can now state that these complexes initiate the process and originate from molecular liquids generated by the secretion of glandular tumor epithelia. These liquids inside the tumor glands generate a clockwise rotating **vortex and counterclockwise** movements, creating real molecular whirlwinds from which the spirals and triangles are formed, the crystalloid proteins of this process, which is in essence an **eminently physical phenomenon**. We wish to draw attention to the images documented below that show how malignant tumor glands adopt an emerging **embryoblast**–**blastocyst-**like function. Remember that the blastocyst is an embryonic structure that forms during the early stages of development in which the morula develops as a fluid-filled cavity, transforming itself into a blastocyst or embryoblast.

In stages 2 to 5 we can see fractal copies of these protein crystalloid entities that have spontaneously replicated.

In stage 6, the entities gather all the information in a single structure and perforations can be seen on its surface. This entity full of holes on its surface and this, which we call Triplet Crystal (Tc) is constituted by a triplet of spirals and pyramidal - triangles perfectly and beautifully assembled spatially, which behaves as a mold-vector of biological information.

#### b) Cellular memory module: Tc crystals

In stages 7 to 10 we can see the b) **Cellular biological phase**, the Tc crystal pattern of stage 6, behaves like a vector of biological information and transfers information to the nearby injured squamous cells, generating a change in the normal polygonal cellular phenotype, and acquiring the same pattern of the pyramidal triangular phenotype of the **Tc.**

In stages 11 and 12 we can see the complete transfer of information from **Tc stage 6 to** squamous cells, generating an exact copy of the Tc phenotype.

We can see how the transformed squamous cells progressively become more hyperchromatic as they reach stage 12, where an embryoblast-like entity is clearly shown. Fractality and chirality are the mediating instruments in this process that complete the metamorphosis experienced by the squamous cells probably regulated by the reactivation of genes and proteins that come from normal embryogenesis.

This evolutionary cycle shows other surprising details that speak for themselves: in the transition phase between stage **10 and 11**, four pairs of **pyramidal-triangular satellite** cells can be seen near the squamous cells that are transforming their phenotype. These are once again perfectly visible in a chiral position (highlighted in a purple circle) in direct correspondence with two (2) embryoid entities in formation that appear one in front of the other with one in a chiral position (these are indicated with a green arrow).

If we link the individual memory modules of protein crystalloid with the spatially separated cells from stages 1 to 12, we obtain the spatial image of a perfect collective memory that has the phenotype of each of the individual memory entities. Here we can see how the collective organization is identical to each of its individual cellular parts and each individual cellular memory has the phenotype of a collective memory encoded.

Fig 1 c, d illustrates the confirmatory biopsy of the cancer in situ as seen in the image, we can verify how the embryo-like patterns that appear on cytology smear are identified in the patient’s biopsy with better resolution and organization. Again 12 sequential steps are observed that recall the blastocyst cavity filled with liquid. In the last stage number 12, the result of this self-assembly order appears, an embryoblast-like - entity with its corresponding chiral mirror image, on the left, clearly bordering with the disorder area. Similar patterns are seen in the cytology smear and biopsy from the same patient, this means that the structure is real and predictable in such a way that we can isolate it.

With the unique pattern identified in this evolutionary cycle, we generated a prototype algorithm that allowed us to perfectly trace these entities in other tumor scenarios, and this is how we were able to identify them in carcinomas, adenocarcinomas, and sarcomas. We had the opportunity to document these patterns in other scenarios under similar conditions.

Figure 2 shows in detail the triplet of modularly integrated components that form the **Tc** entity. Panel a, b, c illustrates **triangular crystals with corresponding pyramidal cells**.

**Figure 2.**
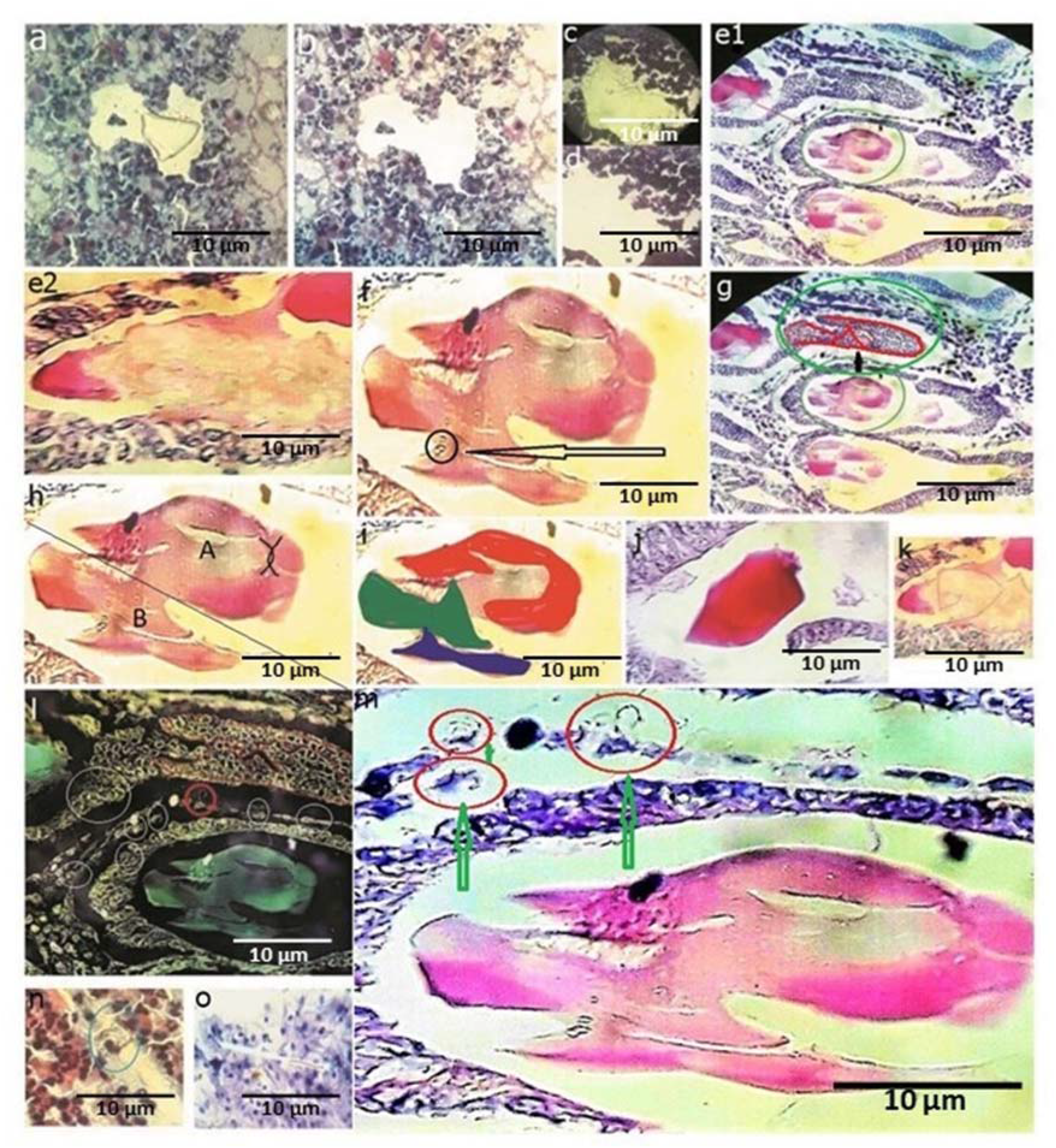
Self-repair order structures: Tc represents the geometric pyramidal triangular spiral complex precursor in the formation of these embryoblast entities. We can observe how the tumor epithelial cells around crystal Tc, have acquired pyramidal cell pattern phenotype with the spiral rotating vortex. Triangular crystal behaves like a vector of biological information and transfers information to the nearby injured glandular cells, generating a change in the normal phenotype as self-repair structure(green arrows) Material was identified in the lumen of a tumor gland in an endometrial adenocarcinoma H. E stain 40X.

We can observe how this entity emerges from the secretory activity of the tumor gland in the lumen of an **endometrial adenocarcinoma** simulating a **blastocyst-embryoblast** (Panels e1-e2). Tc represents the geometric-pyramidal-triangular-spiral cleavage precursor in the formation of embryoblast entities

**The architectural union of three (3) spirals and three (3) crystal triangles organized spatially via rotating vortices in absolute perfection structure is what we call a Triplet Crystal (Tc).** We can observe how the tumor epithelial cells around them, indicated with a red circle, have acquired the Tc **phenotype** (Figure 2 panel m**), generating pyramidal cells.** These images clearly express, as never before, how the geometrical entity that orders Tc can somehow physically regulate the micro-macro cellular environment where these entities are gestated and self-assembled.

The Tc images show how this entity is an interface structure.

In panel h, A shows the **cellular biological** phase of the entity, given by its round phenotype, while B shows protein crystals and a geometric pyramidal triangular-spiral pattern constituting the physical phase of the entity. At present, there is no evidence of a structure produced naturally where this structural conjunction between physics and biology can be so clearly observed, which makes this structure a mold to be copied and used in bioengineering as an entity that generates order and unique biological organization.

Figure 3. panel a illustrates groups of malignant cells aligned in sequences of perfect progressive order, it is reasonable to admit that cells in stage 1 correspond to the totally undifferentiated round immature cells responsible for the highest proliferative growth, these cells are the ones that rapidly metastasize. As the cell advances in these stages, its morphology changes, in stages 4 and 5, as observed, they become pyramidal**–triangular cells** with a spiral component, these cells appear more differentiated in panel b cells are positive for neuro-specific enolase and their potential for tumor growth decreases, material was captured in prostate adenocarcinoma H.E stain 40X. In stage 10, 11, 12 cells have has acquired embryoblast-like morphology organized with a visible head-caudal polarity, these entities enter into symbiosis with the tumor host and are proliferatively indolent and do not metastasize. Patients who express these entities in their tumors have a longer life expectancy than those who do not express it, if we carry out DNA sequencing of stage 10, 11, and 12 cells and compare it with stage 1 DNA sequencing, probably we may find reorganization and repair of mutated DNA segments. This opens up transcendental target therapeutic possibilities for cancer.

**Figure 3.**
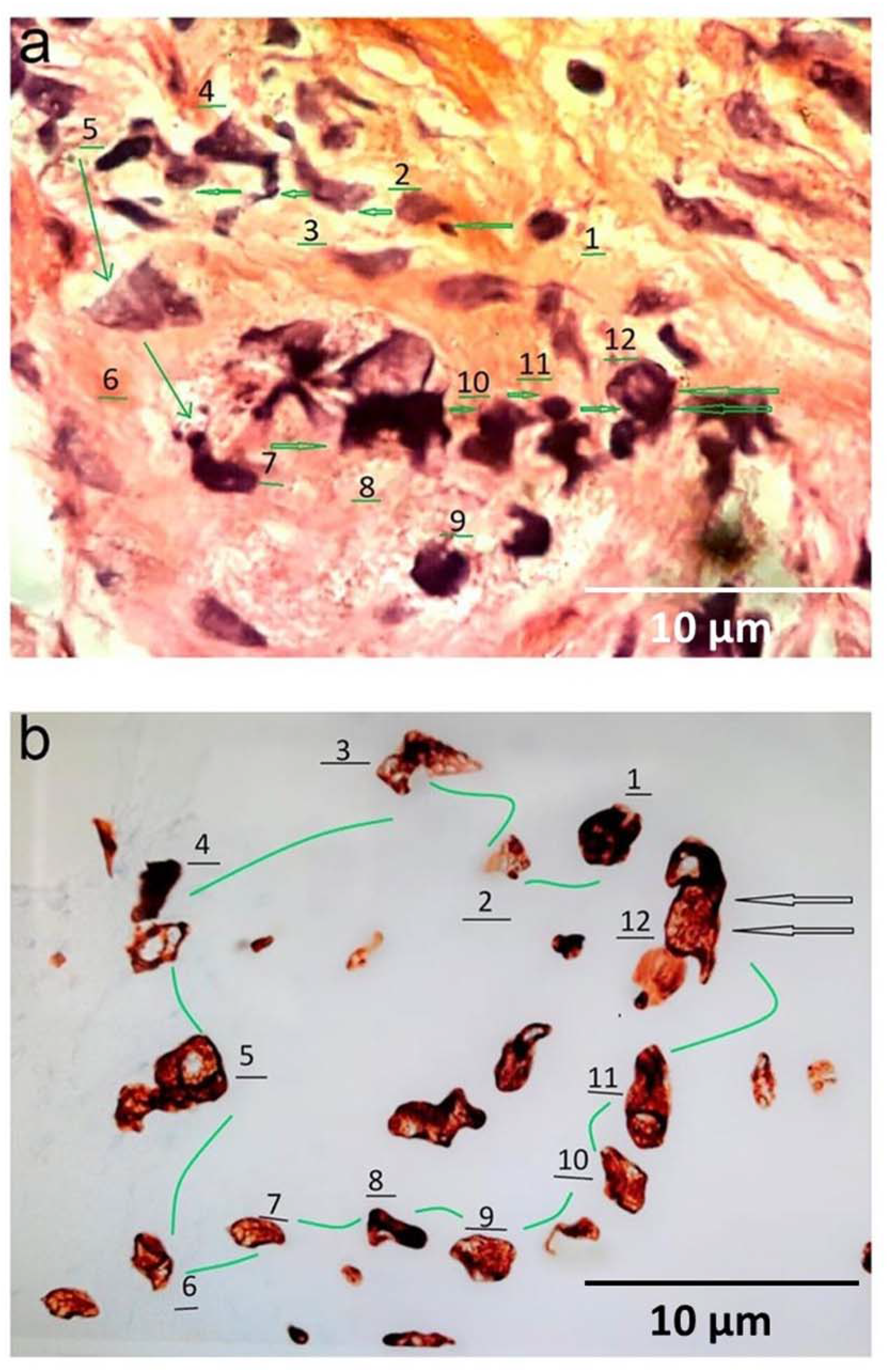
Self-repair structures: Clusters of malignant cells aligned in progressive spatial order sequence via rotating vortices ending in the generation of an embryoblast entity. Material was identified in the lumen of a tumor gland in a prostate adenocarcinoma. H. E stain 40X.

Figure 4 Shows the transfer of information from Tc to a group of tumor cells in several scenarios. Tc represents the geometric pyramidal triangular-spiral cleavage **precursor** in the formation of embryoblast entities. Material was identified in different types of malignant tumor lesions.

**Figure 4.**
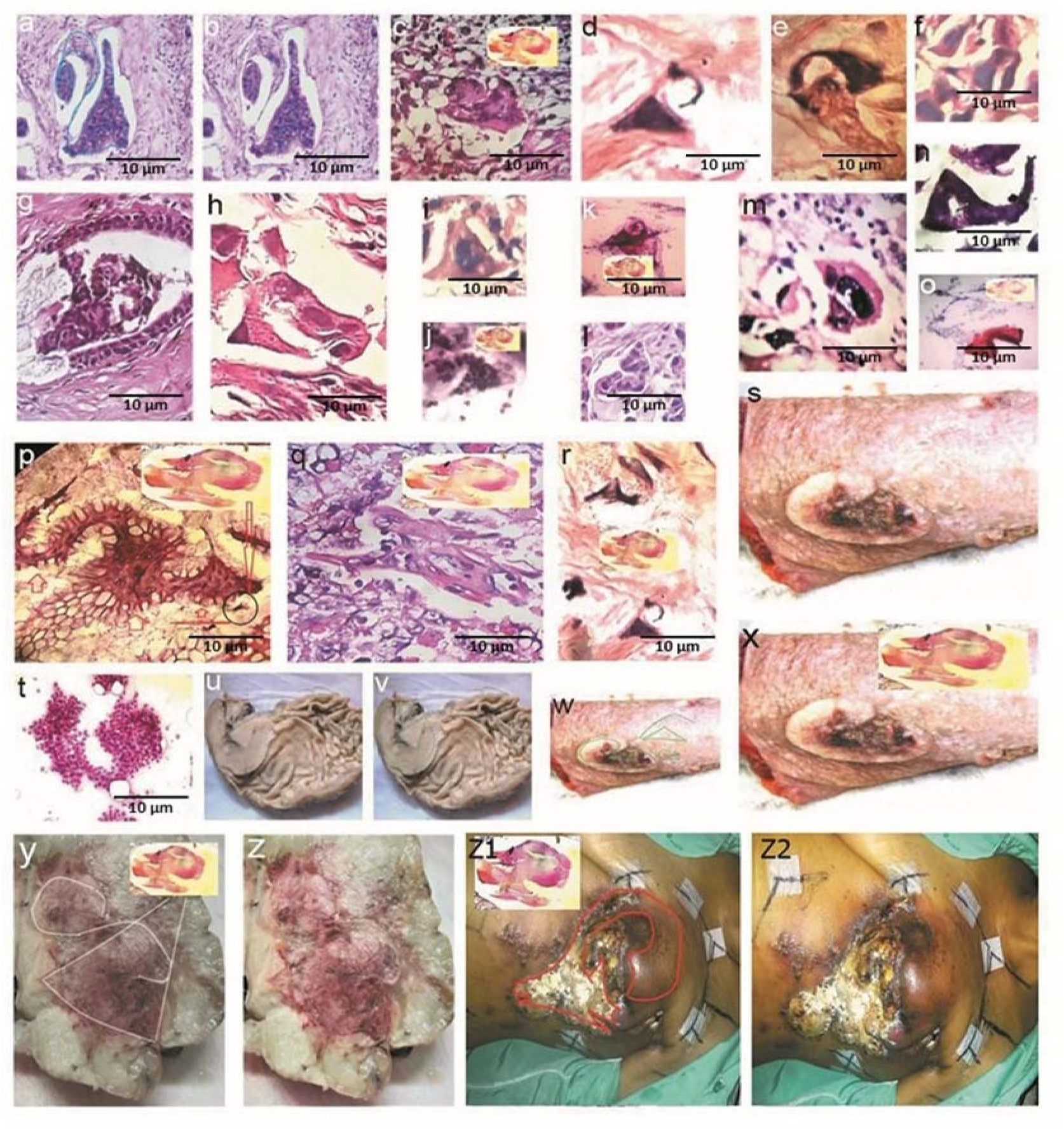
Self-repair structures: Transfer of information from Tc crystal protein to a group of tumor cells in different cancer scenarios. The spiral-pyramidal -triangular rotating vortices organization is illustrated. The “evolutionary state” is represented in stages 8 and 9, the sequence in which Tc transfers information to tumor cells or groups of cells in various scenarios, changing the phenotype of the tumor cells to the crystal phenotype of Tc a, b) breast cancer HE 40 X stain. c) prostate cancer HE 40X stain. d, e, f, g, h) breast cancer HE 40 X stain. i, j, k, l, m, n, o, lung cancer HE 40 X stain. p, q, r) colon cancer HE 40 X stain, s) macroscopic view skin cancer amputation t, u, v) stomach cancer micro -macroscopic view. w, x) macroscopic view skin cancer. y, z) soft tissue sarcoma z1 z2) macroscopic view breast cancer.

Figure 5 shows the final stage where we can observe embryoblast-like memory entities, in different types of cancers which measure from 4 to 12 um, the hallmark of these entities is that they generate **chiral-cell structures** as documented here with the green arrows a) breast adenocarcinoma, illustrate perfect embryoblast structures gestated by the tumor. The green arrows indicate chiral cell mirror images. b) embryoblast-like structure surrounded by necrotic tumor cells. Due to their differentiation, these entities resist cellular hypoxia, evade the immune system, they are probably chemotherapy and radio resistant, and paradoxically, the tumors in which these entities are identified have a better prognosis than those tumors in which they do not occur**. It is logical to assume that these cells and tissues that make up this entity with greater differentiation no longer retain tumorigenic potential and have less invasive growth capacity with pluripotentiality**. c) Sarcoma tumor, the green arrows indicate embryoblast-like entities with chiral mirror images. d.) Osteosarcoma e) cervical cancer cytology smear illustrates perfect embryoblast-like entity with chiral mirror image. f) Brain tumor within rosette-like cavity embryoblast entity with the respective cell chiral image. g) This is one of the most representative images where specific details of organization are recognized and where there is not only cellular but also tissue differentiation with a central vascular tubular axis and cephalo-caudal polarity. In the cephalic pole, the temporal **petromastoid bone and eye periorbital area** is recognized. Material was captured in the ascitic fluid in a patient with colon cancer. h. i) Illustrate NS high immunopositivity of embryoblast-like entities.

**Figure 5.**
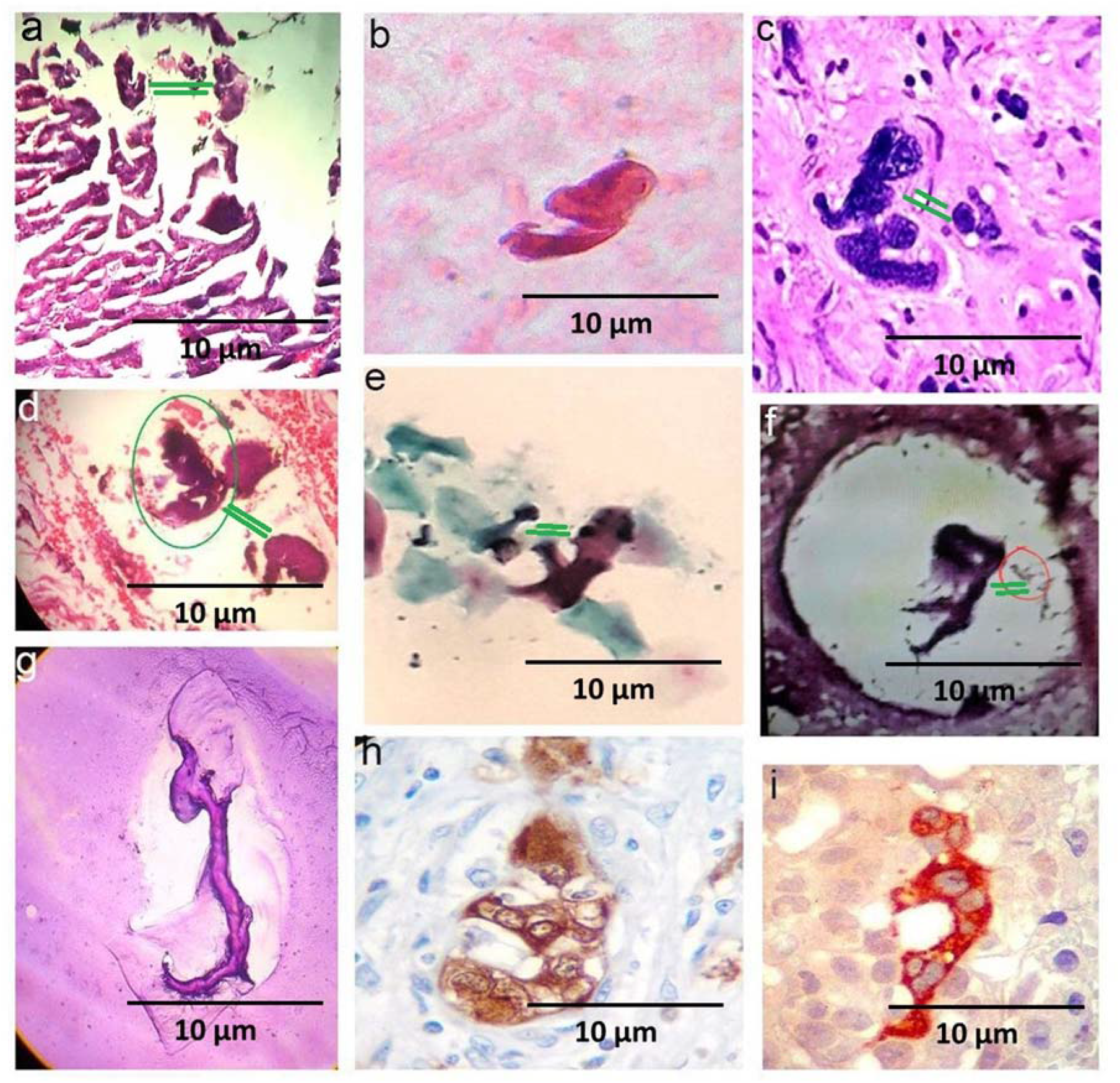
Self-repair structures with pluripotentiality. Cell embryoblast-like entities identified in different types of cancer. Cancer generates self-assembly repair order structures via rotating vortices. Swirlonic state with spatial cephalic-caudal growth organization The hallmark of these entities is that this structure can exist in two copies of visible chiral cell conformations and therefore behaves in accordance with distance molecular communication, which results in clusters of malignant cells aligned in progressive order sequences (green arrows). Panel h, I illustrates perfect embryoblast-like entities with high specific immunopositivity expression for neuron specific enolase,

### Neuron enolase immunostaining

Tc represents the geometric-triangular-spiral cleavage precursor in the formation of embryoblast entities with high immunopositivity expression for neuron specific enolase (95.0%) (table 2). (Figure 3 panel b; figure 5 panel h,)

#### c) Growth memory module

This corresponds to the macroscopic phase of the evolution of these entities that we have documented. To do this, we will rely on specific cases with macroscopic and microscopic histological documentation in this phase of growth..(.Fig 6, Fig 7,Fig 8.)

**Figure 6.**
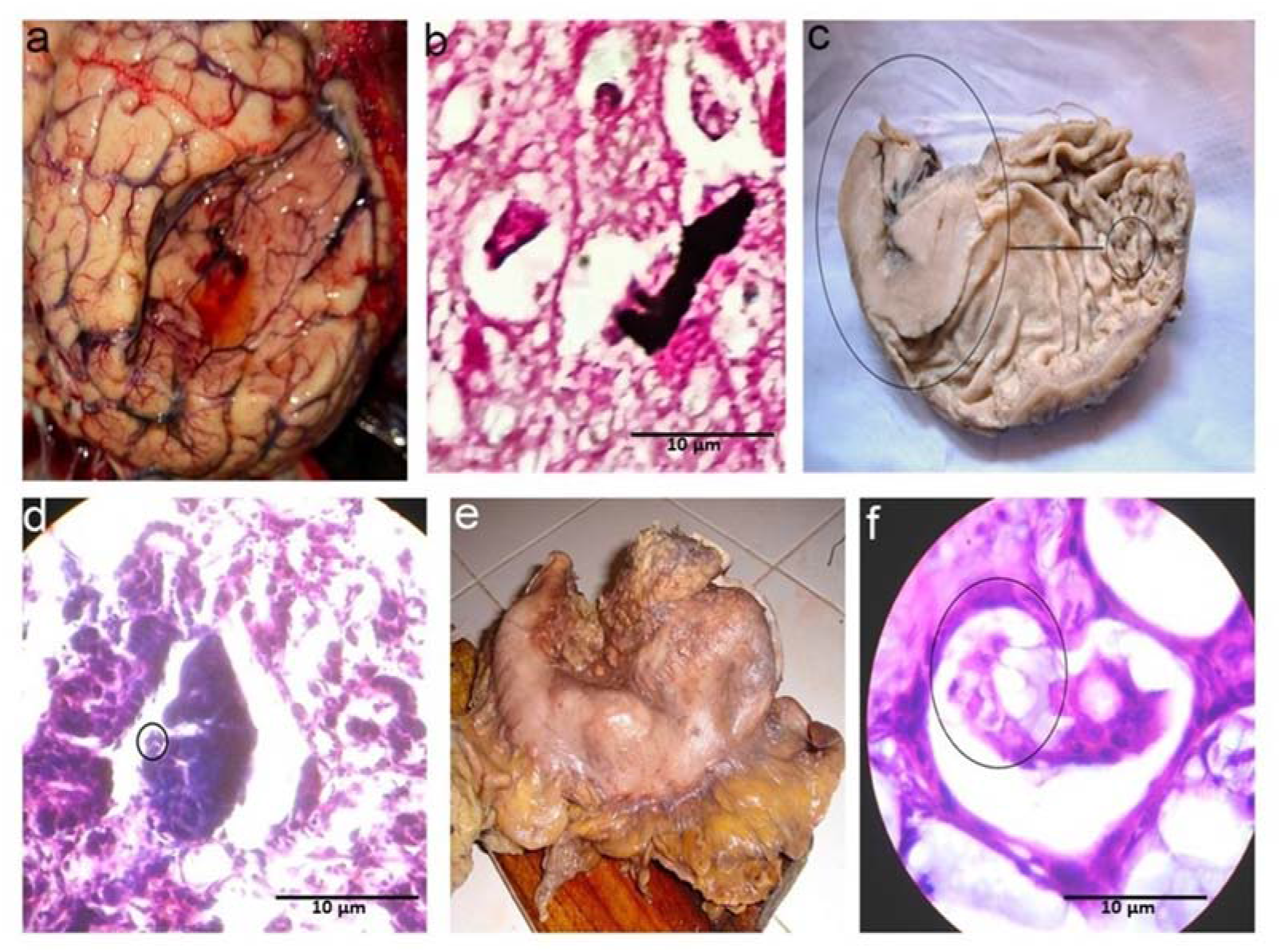
Self-repair structures: Glioblastoma multiforme a) macroscopic phenotype lesion b) microscopic phenotype lesion, it can be seen how the two phenotypes are identical. Panel c, gastric cancer, the microscopic crystalloid phenotype is identical to the macroscopic tissue phenotype. Millions of copies of an individual microscopic crystalloid memory generate a collective macroscopic memory identical to the microscopic one in time (8 to 9 years) through the transfer of information, panel e, f) gastric cancer macroscopic phenotype identical to the microscopic phenotype

**Figure 7.**
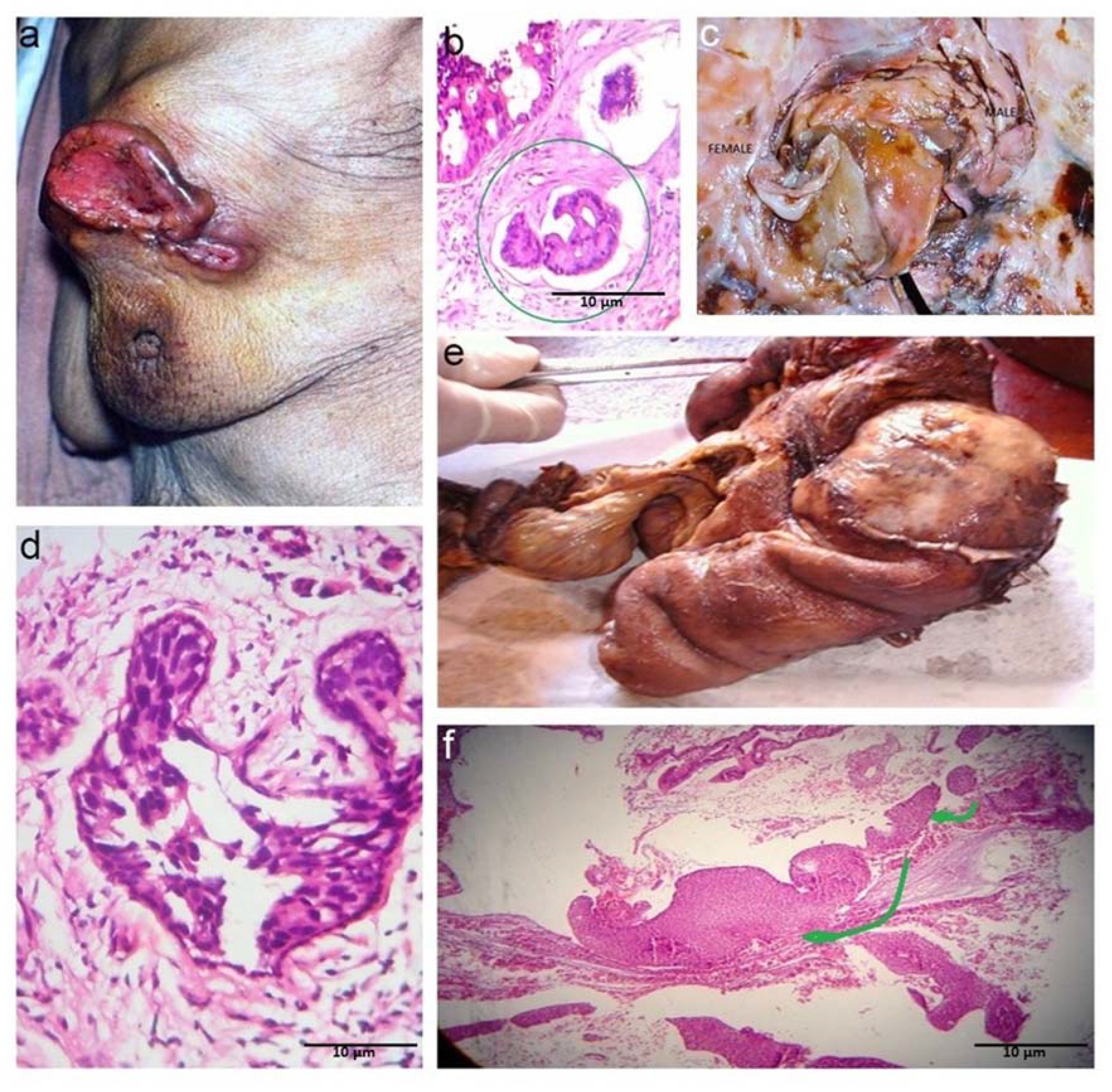
Self-repair structures: a, b. 80-year-old patient with a breast tumor. A giant breast tumor lesion can be seen that clearly shows an unusual morphology of cephalo-caudal pattern with an upper base and a lower base that is also triangular with a bilaminar caudal portion. The microscopic study of this lesion showed an identical lesion with a bilaminar component identical to the one observed macroscopically. The microscopic seed grew in time generating millions and millions of identical fractal copies. c. This is a 50-year-old tumor patient with a uterine leiomyosarcoma, panel c, a nest-like lesion where two (2) triangular structures are seen in a mirror image in chiral position with a spiral component, two (2) structures are seen with caudal-cephalic characteristics; on the left appears an embryoblast XX phenotype pattern due to the curvilinear morphology of its lines, on the right side is another less curvilinear morphology structure with XY phenotype pattern d) illustrates another microscopic embryoblast-like entity with two visible chiral conformations: XY and XX phenotype patterns side to side.. Material was captured in breast adenocarcinoma. H. E stain 40 x. e, f) Small intestine lymphoma, the microscopic phenotype identical to the macroscopic phenotype,

**Figure 8.**
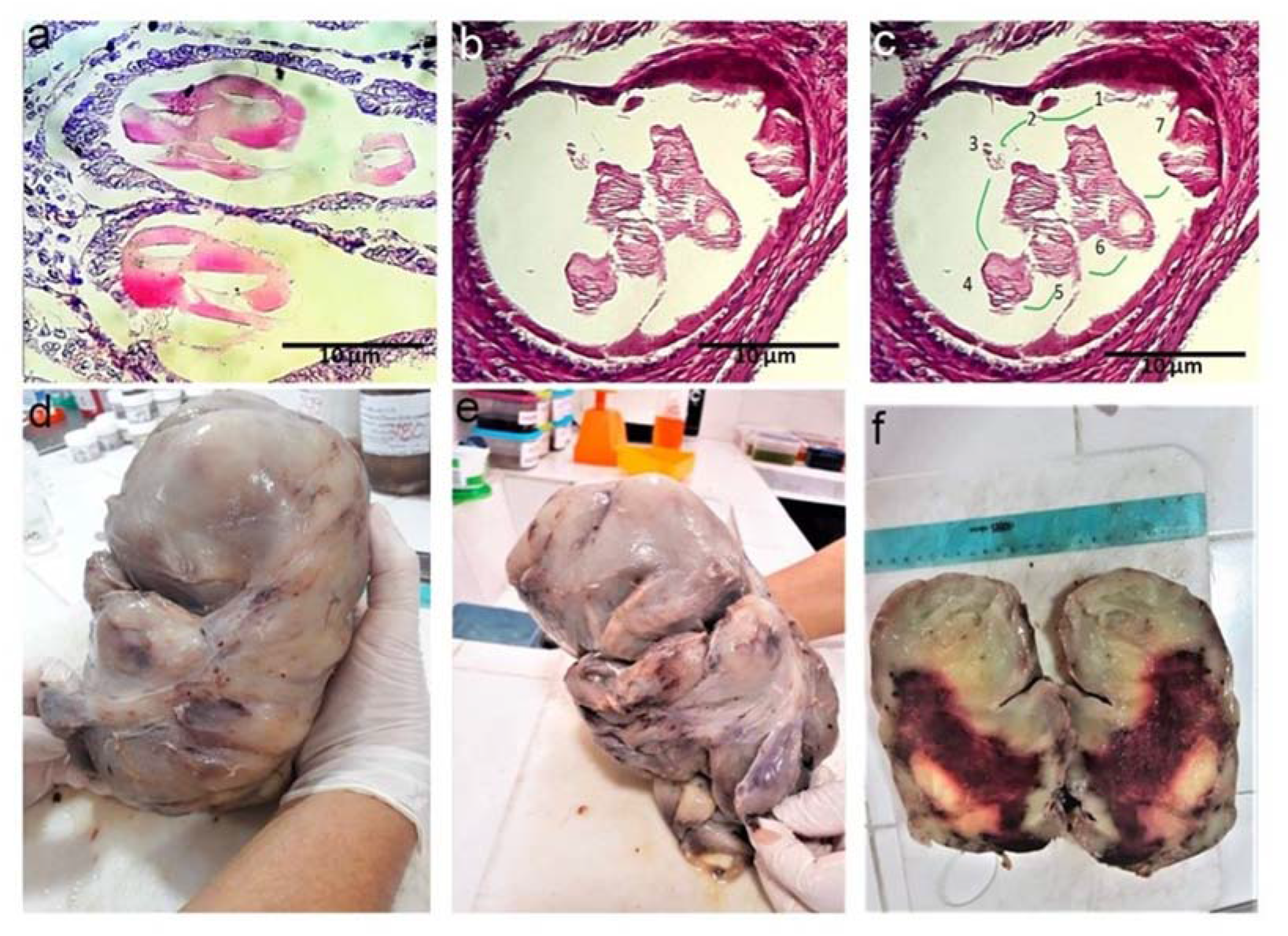
a, b, c, d, e, f) This is a 35-year-old male patient with a retroperitoneal tumor lesion measuring 25 cm in length and weighing 2800 grams. Panels a, b, and c show the perfect tracing that we were able to carry out in this case. Panel a shows how the individual microscopic memory has its genesis inside a tumor gland from a molecular fluid that solidifies and crystallizes. Panel b, c irrefutably illustrate how the fluid secreted by the tumor epithelial cells solidifies and becomes crystalloid in stages 1,2,3,4,5,6,7. From this individual crystalloid memory an easily discernible embryonic phenotype is sequentially formed in stages 3 to 7. Copies of this microscopic crystalloid memory generated an entity with clear fetal-type characteristics over a period of 8 to 9 years, identical to the crystalloid memory phenotype that served as the mold. Panel d, e, shows an almost perfect fetal macroscopic structure in profile and from the front, panel f shows a cross-section of the characteristics of the tumor that was diagnosed as a component of a sigmoid colon carcinoma.

Figures 9 and 10 illustrate the most perfect identified structures in cancer tissues with defined morphological details demonstrating topographical pluripotentiality.

**Figure 9.**
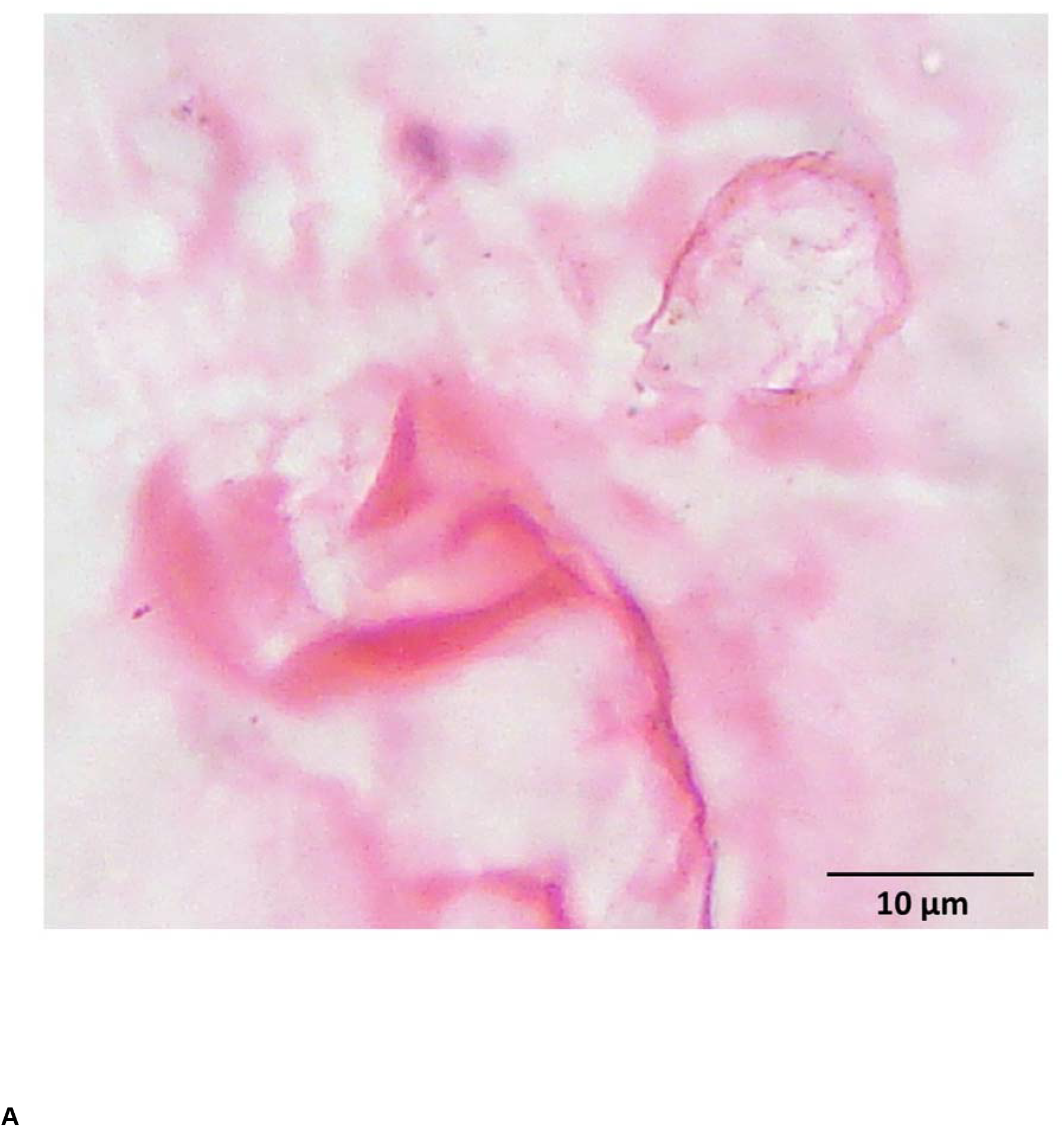
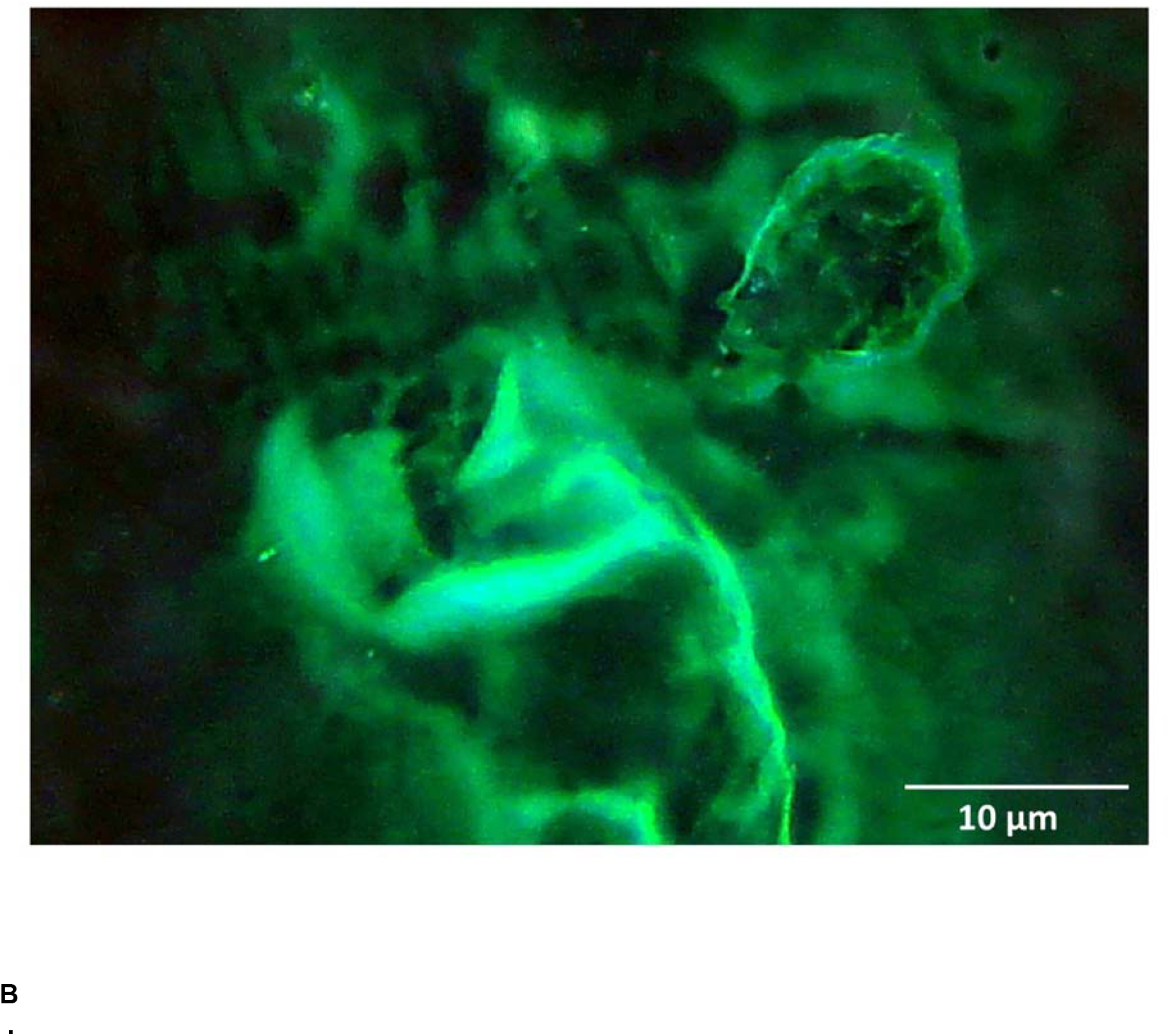
A, B illustrate perfect development of cell embryoblast structure identified in ascitic fluid carcinomatous fluid. A self-repair structure, spatial cephalic-caudal growth organization with pluripotentiality

**Figure 10.**
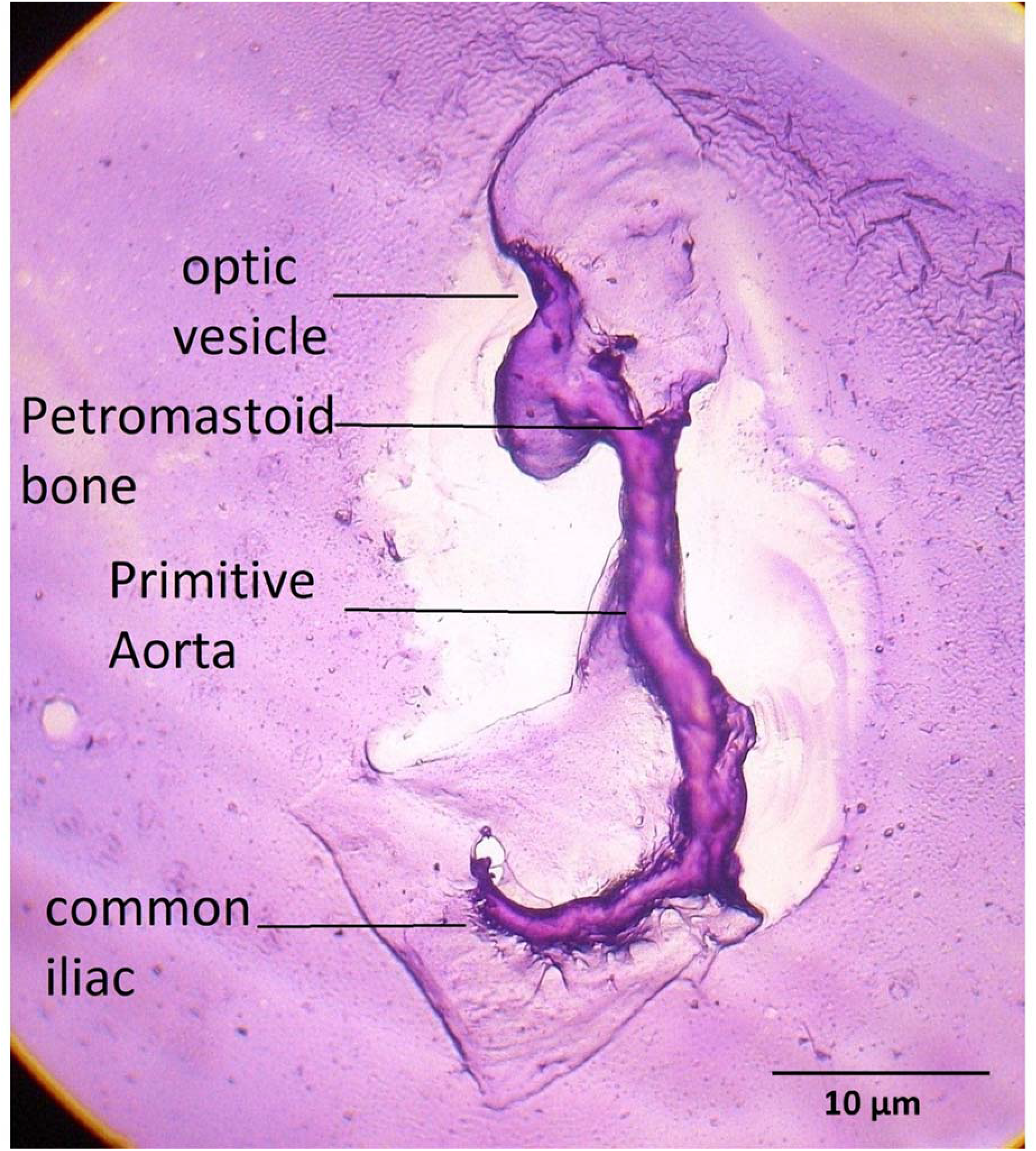
illustrates perfect cell embryoblast structure with defined visible pluripotentiality Observe optic vesicle petromastoid bone primitive aorta and common iliac. Material was captured in ascitic fluid carcinomatosis. A self-repair structure with spatial cephalic-caudal growth organization

## Discussion

We are witnessing a transcendental phenomenon in biology in which evolution patterns and processes shape cancer.

Cancer has two opposite evolutionary signaling pathways: an autonomous proliferative disorder state, monoclonal cell proliferations that generate mass and metastasis that advances at the rhythm of the organism’s chronobiology and ends up collapsing it, and the opposite: a hidden reparative state that reorganizes and generates order returning to the biological past by activating inactive genes of embryogenesis with polyclonality and pluripontentiality. We all know the disorder and death phase …here we document the other side of that coin, the evolutive state that carries the correct DNA instructions to repair and regenerate.

Collective organization of cancer cells can partially or completely return to embryoid genotype-phenotype with the plasticity to transform their morphology in cell embryoblast-like evolutionary memory entities by expression of dormant genes that come from embryogenesis. The hallmark of these entities is that they exist in two copies of visible chiral cell conformations via swirlonic state or rotating vortices, the same one that induces a collective behavior in schools of fish, flock of birds, swarms of insects, which produces in malignant tumors, clusters of fractal cells aligned in progressive order sequences.

After hundreds of driver mutations cancer cells gain new abilities or attributes and recapitulate early stages of the embryogenesis with high immunopositivity expression for neuron specific enolase. Our findings document how malignant tissues reactivated ancestral storage memory and elaborate inside tumor glands spiral-pyramidal -triangular fractal chiral crystals (Tc) as geometric attractors proteins and biomimicry the primitive cellular blastocyst embryoblast fluid-filled cavity. The resultant evolutionary embryoblast-like entity has higher survivability and spatial cephalic -caudal growth organization with pluripotentiality, that carrier the correct DNA instructions to repair and regenerate. The isolation and manipulation of these order structures can guide and control the regenerative pathway mechanism in human tumors as follows : 1) Modify and reprogram the phenotype of the tumor where these entities are generated. 2) Establish a reverse primordial microscopic mold to use the swirlonic collective behavior of cellular building blocks to regenerate injured tissues. 3) Convert cancer cells to a normal phenotype by developmental patterning of active patterning **cues. 4) Convert cancer cells to a normal phenotype by regeneration using the organizational level and scale properties of reverse genetic guidance. 5) Globally control mitotic activity and morphogenetic movements avoiding their spread and metastasis, determining a better life prognosis for patients who incubate these entities in their tumors compared to those who do not express them.**

Below we offer verifiable reasons that show how these entities are really predictable and reproducible entities:

**a)** These entities appear in states of cellular emergency and do not appear in normal tissues

**b)** The hallmark of these entities is that their structure can exist in two copies of visible chiral cell conformations which result in clusters of malignant cells aligned in progressive order sequences via rotating vortices, swirlonic state. fig 5 (green arrows) what involves apical-basal polarization, cell adhesion, cell communication, spatial migration programmed apoptosis and positional signals; Figure 7d illustrates embryoblast-like entity with two visible chiral conformations: XY and XX phenotype patterns side to side

c) Behind the morphogenesis of these embryoblast-like entities appear precursor patterns of pyramidal cell geometry derivate from spiral-triangular crystal geometry **in which Tc crystals change the phenotype of tumor cells**

**c)** The cells that make up these entities show neuroectodermal differentiation and are positive for NSE

**d)** These entities have micro and macroscopic grow representativeness, may show bone andvascular tubule differentiation fig 5g **e)** These structures resist the normal tumor necrosis activity suffered by the tumors, as seen in figure 5b where an embryoblast-like entity is observed in the middle of tumor

**f)** These entities are perfect microscopic versions of a 10-to 13-week-old human embryo

**g)** They emerge within glandular structures or in areas of cystic degeneration and mimicking the fluid-filled blastocyst cavity during embryogenesis

**h) These** entities eliminate all the proliferative power growth from the tumor as they differentiate neuroectodermally, entering into symbiosis with the host preventing metastasis and determining a better prognosis for tumors that express these entities compared to those that do not present them.

The main characteristic of these entities is the identification of clusters of malignant cells aligned in perfect progressive order sequences. Sequence clustering attempt to group biological sequences that are somehow related. constructing a transitive closure of sequences with a similarity. **The similarity is often based on sequence alignment.** These progressive linear sequences remind us of linear sequence of nucleotides along a segment of DNA that provides the coded instructions for synthesis of RNA, which, when translated into protein, leads to the expression of phenotype character. Cells aligned in progressive order sequences must correspond to a molecular guideline of assembly information of genes and DNA. **Knowing the DNA sequence ordering in these entities would have transcendental practical and therapeutic applications**.

These patterns of sequentiality and order show that in the midst of the biological chaos and mutations that cancer represents, there is a product in development that is “gestating” as a result of the reactivation of signals from genes and proteins that return from embryogenesis. **This also makes us think in a reciprocal way that in the initial stage of embryogenesis rotating vortices and swirlonic state can play an important role in morphogenesis.**

We are likely facing an emergent evolutionary biological response, we postulated that these entities that we are documenting are not there randomly, they have a function with the capacity to repair, modify, and/or regenerate damaged senescent and mutated cells.

The **genesis** of these entities is given by the **fluid** secretions within the tumor glands that rotate in opposite **vortices**, and solidify forming crystals **of spiral pyramidal triangular** geometry, representing the crystalloid structural protein of these entities, which measure just a few microns but acquire polarity, organize, fuse and generate a triplet of triangular images and spirals forming a conglomerate that we have called triplet crystal (Tc). **Tc represents the geometric pyramidal-spiral cleavage precursor in the formation of these embryoblast-like entities**.

In articles such as *Magnetization of the three-spin triangular Ising model*, theoretical physicists have already used mathematical formulas to address the organizing power of these geometric conglomerates that take shape now in cancer biology and become real in practice in these self-organizing entities that we have documented here for the first time. **Tc** is a new, unique structure that nature builds at the interface of physics and biology where **Geometric swirlonic systems hold promise for finding new phases of biological self-assembly**. [16 17]

The collective cancer system cell organization probably start with self-propelled particles (SPP) that interact with each other, which can lead to the emergence of **collective** behaviors. These collective behaviors mimic the self-organization observed with the flocking of birds, the swarming of bugs, the formation of herds of sheep.

In 2020, researchers from the University of Leicester reported a hitherto unrecognized state of self-propelled particles — which they called a “**swirlonic state**” [18] The swirlonic state consists of “swirlons”, formed by groups of self-propelled particles orbiting a common center of mass. These quasi-particles demonstrate a surprising behavior: in response to an external load, they move with a constant velocity proportional to the applied force, just as objects in viscous media do. Swirlons attract each other and coalesce forming a larger, joint swirlon. Typical collective motion generally includes the formation of self-assembled structures, such as clusters and organized assemblies. The prominent and most spectacular emergent large-scale behavior observed in assemblies of SPP is directed collective motion. In that case all particles move in the same direction. On top of that, spatial structures such as bands, vortices, asters, and moving clusters can emerge. One of the key predictions of the SPP model is that as the population density of a group increases, an abrupt transition occurs from individuals moving in relatively disordered and independent ways within the group to the group moving as a highly aligned whole**. Thus, in the case of cells in malignant tumors, this creates a trigger point in which disorganized and dispersed locusts transform into a coordinated marching army. When the critical malignant cell population density is reached, the tumor cells should start marching together in a stable way and in the same direction generating spatial self-assembly spiral-pyramidal-geometric order**

The theory of physics of a magnetization model forming a perfect conglomerate of a three-spin, three-triangle as is shown in the figures of the paper. In detail, a slender crystal-like Tc is able to behave as a vector of biological information and transmit its phenotype to squamous cells that change their phenotype in response to this transmitted information. As we can clearly see in the “evolutionary cycle” these cells slowly and progressively change their phenotype, turning into a final product; cell embryoblast-like memory entities. This visible transfer of information from Tc to the tumor cells reminds us of the hypothesis of Professor Cairns-Smith, A. G when he states that crystals can behave as genes mainly in relation to cancer. [19]

What are these entities and what do they represent? Cell embryoblast**-like memory entities determine how the traces of the initial phase of development of blastocyst-embryoblast are indelibly engraved on and stored forever in the individual and collective memory of all the normal cells of the human body** which we believe represent the genetic basis for tissue regeneration and repair, as mainly observed in the glandular epithelium. This memory is activated in a state of cellular emergency, disorganization, or cellular senescence. Perfect microscopic replicas of morphological information that possess the information of a collective memory encoded in its nucleus and anchored to the period of embryogenesis. Every cell that is irreversibly injured and escapes programmed cell death or apoptosis has the plasticity of metamorphosis and the ability to return to an embryonic stage. This translates into a transcendental biological fact: Every cancer cell returns partially or totally to an embryonic genotype-phenotype

This article is the consolidation of over ten (10) years of work [2, 21]. **A recent publication in an animal model by researchers from the neuroscience department of the University of California support our findings in human tissues, as they found that adult neuronal cells in mice exposed to cell damage return to a state of embryonic transcriptional growth state [[22].** These findings are of unquestionable crucial significance for our research, which determines that the methodology we used and the data we collected are reliable to the point that they can be reproduced and predicted in other laboratories around the world.

The hypothesis linking cancer and cell damage to embryogenesis is not new. Cohnheim suggested in 1882 [23] that tumor cells were essentially “embryonic” in nature, being remnants of embryonic epithelial cells. In the early 1970s, Brinster [24] demonstrated that by injecting embryonic carcinoma cells into a mouse blastocyst, the mouse was able to regulate the cancer cells and their progeny to the point that they no longer behaved malignantly; rather, they participated in normal embryonic development that resulted in functional mice. This experiment was confirmed by Mintz and Illmensee [25] and Papaioannou [26]. Pierce [27, 28], showed that this effect, specific for some types of tumor cells, is strongly position dependent: the carcinoma cells placed between the pellucid zone and the trophectoderm (the perivitelline space) were not controlled, while the carcinoma cells injected into the blastocele lost their tumorigenicity immediately after differentiation. Clinical trials, conducted with zebrafish embryo extracts administered to patients with advanced cancer that did not respond to conventional treatments, significantly reduced the expression of oncofetal antigens (such as AFP) [29] and induced marked beneficial effects (induction of objective responses, improvement in state performance and significant increase in overall survival) [30 -31].

Additionally, our morphological findings fit perfectly with the new hypothesis by molecular biologist Jose A from the University of Maryland who states that DNA is only “the list of ingredients” and not the set of instructions used to build and maintain a living organism. These instructions are very complex and are stored inside each individual cell as a shape memory that “decides” how and to what extent to use the ingredients available in the DNA. “DNA cannot be seen as the ‘blueprint’ for life” [33, 34].

According to Dr. Jose the fundamental aspects of anatomy are dictated by something outside of DNA and he proposes that non-coding instructions in DNA are actually contained in the architectural arrangement of molecules within cells and in the interactions between them. This arrangement (shape memory) is what is preserved and transmitted from one generation to the next. The findings of our study agree with the observations of Jose *et al* from the perspective that the entities found have their own identity and are unique since they have the differential phenotype of the host where they were gestated. In addition, the presence of visible pores or perforations also reported in the basement membrane of mouse embryos [Kyprianou et al [35] is noteworthy, supporting the theory that the entities found by us present characteristics of real “microscopic” embryos.

The analysis of living systems from molecular to population scales has revealed how the storage and processing of information across multiple scales is a key attribute of life, in which order can arise through the spontaneous association of molecules in the living system and the formation of dynamic structures (Tc) that can store and retrieve information from collections of self-assembled and self-organized molecules gestated in a swirlonic state. These different ways of changing entities, sensors, and properties highlight the multi-scale nature of living systems and suggest the usefulness of different entity-sensor-property frameworks at different molecular-microscopic-macroscopic scales that cannot be explained solely from the perspective of DNA or genome analysis.

## Conclusions

Our findings document how malignant tissues reactivated ancestral storage memory and elaborate; crystalline proteins repaired copies of the damaged substrate tissue. The resultant evolutionary embryoblast-like entity has higher survivability and spatial cephalic-caudal growth organization with pluripotentiality that carries the correct DNA instructions to repair, isolate, and manipulate, these structures can guide and control the regenerative pathway mechanism in human tumors

These observations confirm the importance of the fetal characteristics of cancer as a source of useful information that can contribute to better understanding of the biology of cancer at molecular, cellular, and micro-environmental levels.

These structures clearly express, as never before, how a geometrical swirlonic entity (Tc) can somehow physically regulate the micro-macro cellular environment where these entities are gestated. We can predict these structures based on the knowledge of microscopic forces of self-assembly. At present, there is no documented structure produced naturally, where this structural conjunction between physics and biology is so clear and perfect, which makes it a real platform to be copied and used in the assembly and design of new proteins and artificial biostructures.

Finally, we believe that these evolutive collective cellular embryoblast entities are ready to be removed from their tumor microenvironment, as a self**-repair structure** with pluripotentiality. **An emergent self-repair structure, biological template to develop targeted therapeutic alternatives not only in cancer but also in the treatment of autoimmune, viral diseases, and in regenerative medicine and rejuvenation.**

**Graph 1.**
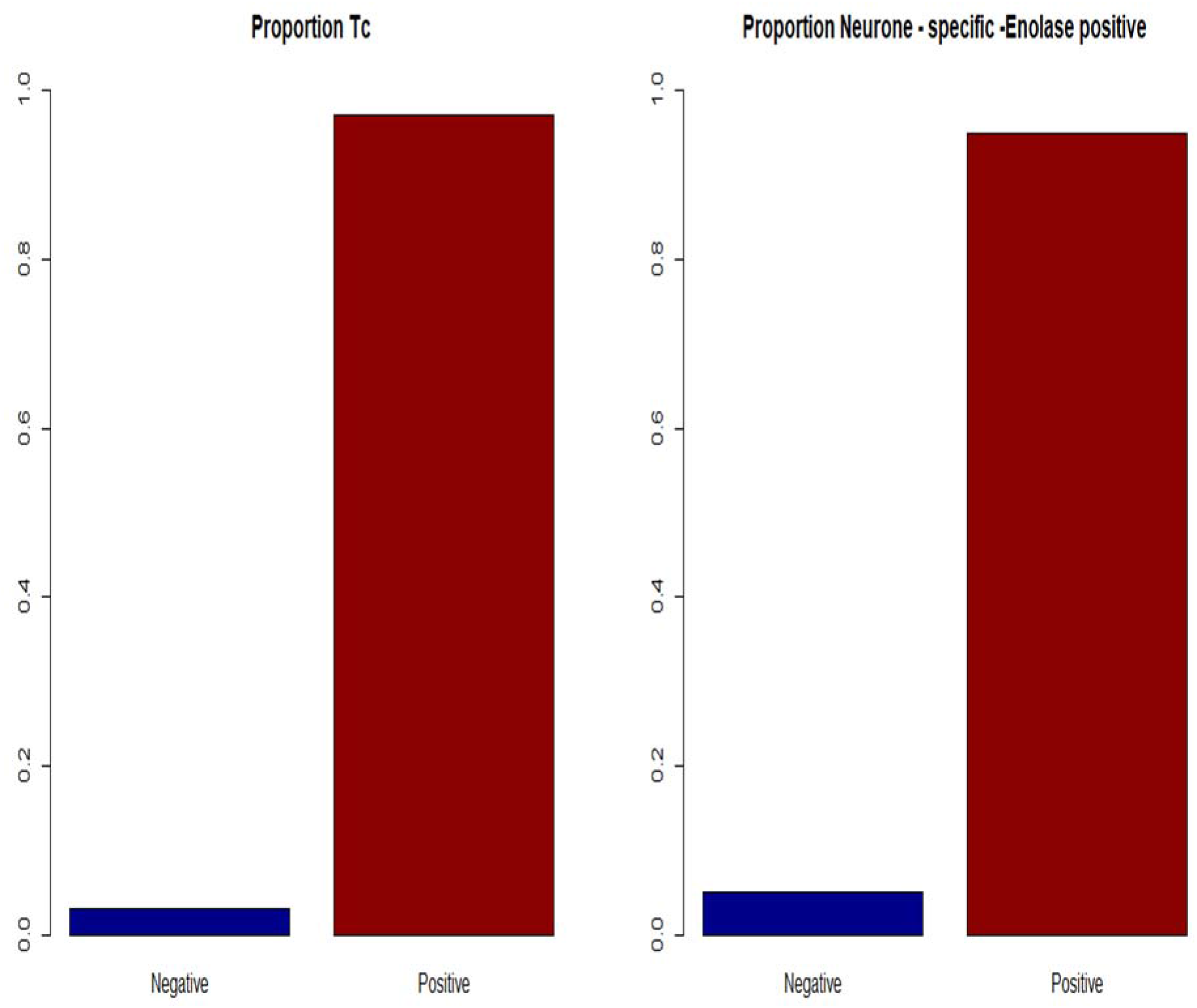
Prevalence proportion Tc and Enolase immunostaining.

## Abbreviations

GTCHC: Geometric triangular chiral complexes
NSE: Neuron specific enolase
Tc: Triplet crystal

## Declarations

### Ethics approval and consent to participate

This study was approved by the ethics subcommittee of the University Cooperative of Colombia, Villavicencio, Colombia, and followed the guidelines of the Ministry of Health (No. 8430 of 1993) and the principles established by the Declaration of Helsinki. All patients signed an informed consent form for the use of their biological materials for diagnostic and research purposes. **Participants consent to publish this information maintaining anonymity.**

### Availability of data and materials

The data sets analyzed during the current study are available from the corresponding author, Competing interests

The authors declare that they have no conflicts of interest.

### Funding

**This study was carried out with the authors’ own resources. There was no external funding**. Authors’ contributions

J.A.D., L.S, and L. A. D guided the project, wrote the paper, and analyzed the results. M.F.M L.C.P K.T.M., O. F. S., M. A.C. and L.K.S recollected and processed the samples**. All authors have read and approved the manuscript.**

## Acknowledgements

The authors thank the board of directors and authorities of the Departmental Hospital of Villavicencio Meta and Dr Jesus Emilio Rosado, director of the Departmental Hospital of Granada Meta) for the logistical support in the realization of this project.

## Notes

Declaration of conflict of interest: None.

### Competing Interest Statement

The authors have declared no competing interest.

### Summary of Updates

we incorporate new images,,tables we modificated the abstract

